# A novel Ca^2+^ signalling pathway co-ordinates environmental phosphorus sensing and nitrogen metabolism in marine diatoms

**DOI:** 10.1101/2020.05.13.090282

**Authors:** K. E. Helliwell, E. Harrison, J. Christie-Oleza, A. P. Rees, J. Downe, M. M. Aguilo-Ferretjans, L. Al-Moosawi, C. Brownlee, G. L. Wheeler

**Affiliations:** Marine Biological Association, The Laboratory, Citadel Hill, Plymouth. PL1 2PB; Biosciences, College of Life and Environmental Sciences, University of Exeter, Exeter, EX4 4QD UK; School of Life Sciences, University of Warwick, CV4 7AL, UK; Plymouth Marine Laboratory, Plymouth, Devon PL1 3DH, UK; School of Ocean and Earth Science, University of Southampton, Southampton, SO14 3ZH, UK

## Abstract

Diatoms are a diverse and globally important phytoplankton group, responsible for an estimated 20% of carbon fixation on Earth. They frequently form spatially extensive phytoplankton blooms, responding rapidly to increased availability of nutrients including phosphorus and nitrogen. Although it is well established that diatoms are common first-responders to nutrient influxes in aquatic ecosystems, little is known of the sensory mechanisms that they employ for nutrient perception. Here we show that diatoms use a novel and highly-sensitive Ca^2+^-dependent signalling pathway, not previously described in eukaryotes, to sense and respond to the critical macronutrient phosphorus. We demonstrate that phosphorus-Ca^2+^ signalling is essential for regulating diatom recovery from phosphorus limitation, by controlling rapid and substantial increases in nitrogen assimilation. Phosphorus-Ca^2+^ signalling thus mediates fundamental cross-talk between the vital nutrients P and N to maximise resource competition, and likely governs the success of diatoms as major bloom formers in regions of pulsed nutrient supply. Importantly, our study demonstrates that distinct mechanisms for nutrient sensing have evolved in photosynthetic eukaryotes.

## Introduction

Marine phytoplankton contribute almost half of global primary production, and are a major sink for rising atmospheric CO_2_ (Field et al., 1998). Diatoms are a critically important phytoplankton group, accounting for ∼50% of organic carbon exported to the ocean interior (Dugdale and Wilkerson, 1998). Evolutionarily distinct from plants and metazoans, diatoms belong to the stramenopile lineage, a major eukaryotic group that also includes plant pathogens (such as oomycetes), flagellates, and brown macroalgae. A key attribute contributing to the environmental significance of diatoms, is their ability to form spatially extensive algal blooms (Buchan et al., 2014). Diatoms frequently dominate the primary phase of spring blooms, outcompeting other phytoplankton taxa by rapidly responding to environmental cues, including increased nutrient availability (Buchan et al., 2014). In coastal systems, where diatoms thrive, nutrient supply can vary dramatically over diverse spatiotemporal scales (Jordan and Joint, 1998; Kämpf and Chapman, 2016; Litchman, 2007). The ability of diatoms to dominate phytoplankton assemblages in such regions of pulsed nutrient supply suggests that they possess sophisticated mechanisms for nutrient sensing. However, the sensory mechanisms enabling diatoms to rapidly respond to nutrient resupply remain poorly understood.

Phosphorus is a major factor controlling ocean productivity (Dyhrman et al., 2007). Limitation by this nutrient is documented in a variety of marine environments (Thingstad et al., 1998; Wu et al., 2000), including coastal ecosystems where co-limitation of diatom populations by phosphorus and Si has been reported (Ly et al., 2014). This has been exacerbated by anthropogenic activities causing shifts from nitrogen to phosphorus limitation in certain coastal waters (Burson et al., 2016). Certainly, bloom simulation experiments have demonstrated the importance of phosphate in controlling bloom dynamics (Alexander et al., 2015; Wurch et al., 2019). Additionally, in highly-productive photic benthic biofilms, the distribution of phosphate can be patchy (Paytan and McLaughlin, 2007). The selective chemotaxis of diatoms towards phosphate (but not nitrate) (Bondoc et al., 2019), suggests phosphate may thus be an important driver of biofilm community structure too.

Diatoms show numerous adaptive strategies for coping with phosphorus limitation. Upregulation of phosphate transporters is well-documented (Cruz de Carvalho et al., 2016). Moreover, enhanced expression of alkaline phosphatases and/or phosphodiesterases, increases phosphorus scavenging capacity (Alipanah et al., 2018; Cruz de Carvalho et al., 2016; Dyhrman, 2016; Dyhrman et al., 2012; Lin et al., 2013; Yang et al., 2014). Diatoms, and other phytoplankton, also substitute phospholipids with non-phosphorus forms to decrease cellular demand (Martin et al., 2011; Van Mooy et al., 2009). A transcriptional regulator, distantly related to phosphate starvation regulator protein (PSR1) of *Chlamydomonas* (Wykoff et al., 1999), was recently found to coordinate such metabolic adaptations in diatoms (Sharma et al., 2019). However, these studies primarily focus on mechanisms underpinning phosphorus limitation responses. Comparatively little is known about the short-term recovery responses of phosphorus-stressed diatom cells to resupply and how they are regulated. Certainly, lipid remodelling occurs within just one cell division following phosphate amendment in *Thalassiosira* (Martin et al., 2011). Yet, understanding of the sensory systems coordinating rapid cellular recovery to newly available phosphate in diatoms, and other eukaryotic phytoplankton, are completely unknown. As these mechanisms likely underpin competitive bloom dynamics, particularly in regions of pulsed nutrient supply, this represents a major knowledge gap.

New insights into nutrient perception mechanisms in other eukaryotes are emerging. Vascular plants use the versatile second messenger Ca^2+^ for sensing nitrate (Liu et al., 2017; Riveras et al., 2015), and Zn^2+^ (Behera et al., 2017), but not phosphate (Matthus et al., 2019). Nitrate resupply to nitrogen-limited *Arabidopsis* plants induces [Ca^2+^]_cyt_ elevations, which triggers several nitrate-associated regulatory responses, orchestrated via Ca^2+^-dependent protein kinases (Liu et al., 2017). This work raises important questions about the role of Ca^2+^ signalling in nutrient sensing in eukaryotes more broadly. Certainly, diatoms use Ca^2+^ signalling for perception of several abiotic and biotic stimuli (Falciatore et al., 2000; Helliwell et al., 2019; Vardi et al., 2006). Moreover, our recent identification of a novel class of voltage-gated channels in diatoms (EukCatAs), demonstrates that they have evolved unique mechanisms for environmental perception in the oceans (Helliwell et al., 2019). Here, we report the discovery of a phosphorus-Ca^2+^-signalling pathway that is essential for phosphorus sensing and acclimation in *Phaeodactylum tricornutum*, and likely provides a competitive advantage to diatoms in regions of pulsed or intermittent phosphorus supply.

## Results

### Discovery of a novel phosphorus-Ca^2+^ signalling mechanism for sensing phosphorus resupply

To investigate the role of Ca^2+^ signalling in nutrient sensing in diatoms, we used a transgenic *Phaeodactylum tricornutum* line (PtR1), encoding the genetically-encoded fluorescent Ca^2+^ biosensor, R-GECO1 (Helliwell et al., 2019; Zhao et al., 2011). PtR1 cells were grown in f/2 medium (Guillard and Ryther, 1962) made up in natural seawater (NSW), but with reduced concentrations of phosphate, nitrate or f/2 trace metals (Materials and Methods). We then monitored single-cell R-GECO1 fluorescence of nutrient deplete cells, following resupply with each respective nutrient. We observed that cells grown in phosphate-limited conditions (1.8 µM) for four days exhibited rapid, transient elevations in cytosolic Ca^2+^ following perfusion with seawater containing phosphate restored to 36 µM (**Figure 1a and b; Video S1**). No such response was detected in phosphate-replete cells. Nor did we detect Ca^2+^ elevations in cells grown with limiting nitrate, or f/2 metals, following resupply with these nutrients (**Figure S1a**). These data suggest that a specific Ca^2+^-signalling pathway, which is activated only under phosphorus limitation, is involved in regulating rapid cellular acclimation to phosphate resupply. By comparison, we found no evidence for a role for Ca^2+^ signalling in sensing nitrate (or trace metals), which is distinct from what has been observed in plants (Liu et al., 2017).

As only P limited cells exhibited [Ca^2+^]_cyt_ elevations following phosphate resupply, we wanted to examine further the relationship between P depletion and phosphate-Ca^2+^ signalling. We grew PtR1 cells in different phosphate regimes: i) phosphate-replete (P_replete_, 36 µM), ii) phosphate-limited (P_limited_, 1.8 µM), or iii) no phosphate amendment (P_0_) over eight days (**Figure S1b**). We observed that exogenous phosphate concentrations in the medium reduced from 1.8 µM to 0.1 µM within just 2 days in P_limited_ cells (initial concentrations in P_0_ (NSW) medium were already very low, at 0.2 µM) (**Figure 1c, inset**). Furthermore, growth of cells in P_0_ and P_limited_ treatments was significantly impaired compared to P_replete_ conditions after 3 and 4 days, respectively (**Figure S1b**). Similarly, Fv/Fm (a proxy measurement of photosynthesis (Maxwell and Johnson, 2000)) values were also reduced in the low P treatments (**Figure S1c**). Phosphate resupply experiments at different time-points revealed that after just one day of growth in P_0_ conditions, cells exhibited the phosphate-Ca^2+^ signalling response following phosphate resupply (**Figure 1c**). Maximal amplitude of the response was exhibited on day 2, and gradually decreased at subsequent time-points. By comparison P_limited_ cells exhibited the response after four days, when cell division slowed (**Figures S1b**). We did not detect phosphate-Ca^2+^ signalling in P_replete_ cells, at any of the time points. Thus, only *P. tricornutum* cells experiencing phosphate limitation exhibit phosphate-induced Ca^2+^-signalling responses.

**Figure 1.**
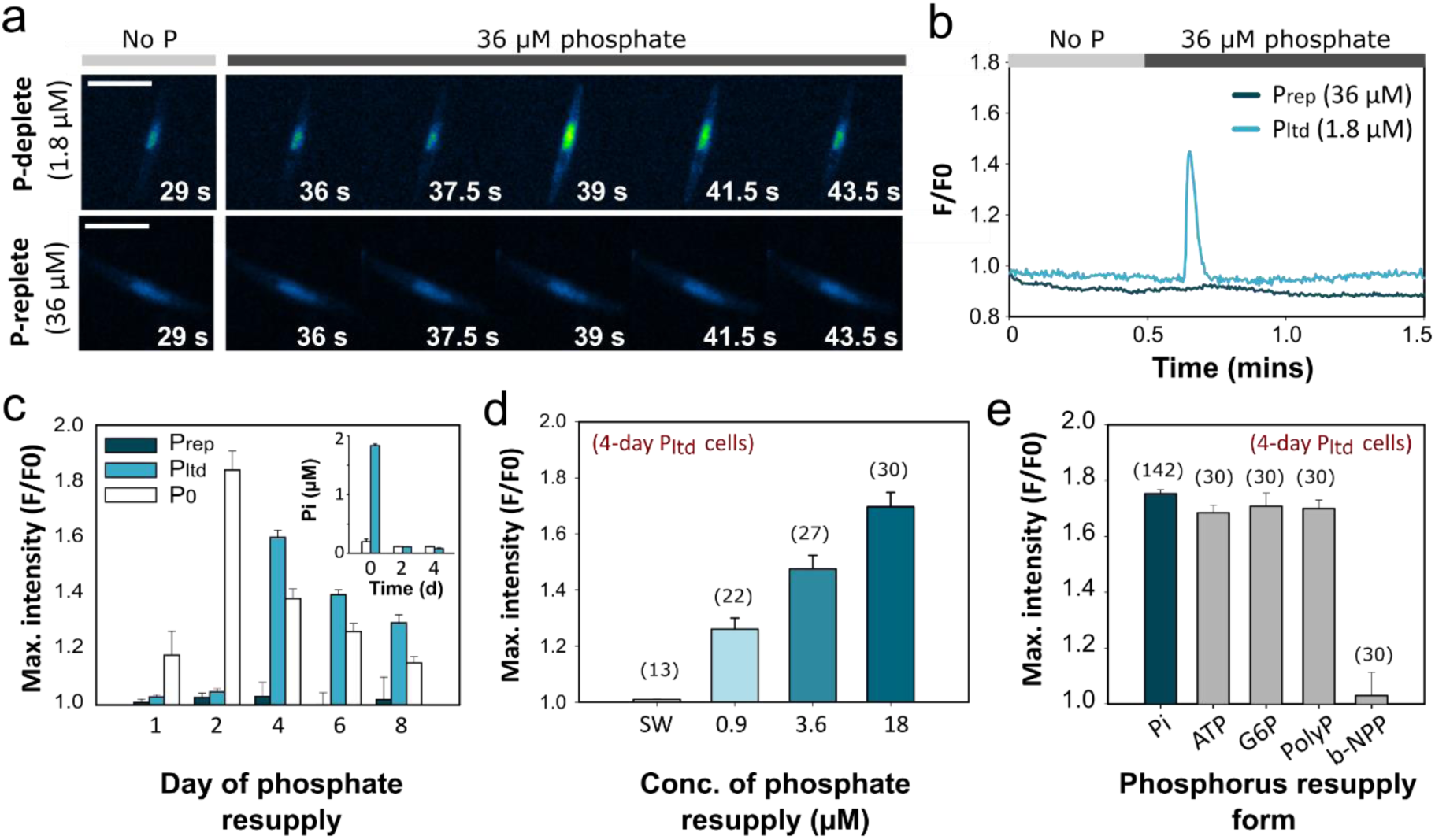
Increases in environmental phosphorus levels trigger rapid [Ca^2+^]_cyt_ elevations in phosphorus limited *Phaeodactylum tricornutum* cells. **a)** Time-lapse images of PtR1 *P. tricornutum* cells grown for four days in f/2 medium in natural seawater (NSW) in either P-deplete (1.8 µM), or P-replete (36 µM) conditions, following resupply with phosphate (36 µM). Cells were pre-perfused with standard NSW f/2 medium without phosphate for 30 secs prior to perfusion with f/2 medium (including 36 µM phosphate). Time stamps indicate the time (s) from the beginning of the perfusion experiment, scale bar: 10 µm. An image of the cell just prior to phosphate resupply (i.e. at 29 s) is shown on the left. The initial blue signal represents chloroplast auto-fluorescence. The experiment was conducted at least three times on independent samples of cells, with similar results. **b**) Representative fluorescence traces (F/F_0_) of PtR1 cells for the experiment shown in (a). **c**) Mean (± SEM) maximal fluorescence (F/F_0_) of PtR1 cells grown in different concentrations of phosphate over 8 days (including P_replete_, P_limited_ and P_0_ treatments with 36 µM, 1.8 µM and 0 µM of phosphate for each treatment, respectively), following phosphate resupply (with 36 µM). Three independent replicates were set up per treatment, with a sample of *n* ≥ 6 cells examined per independent replicate. Inset displays the concentration of phosphate measured in the external medium for P_0_ and P_limited_ cells after 0, 2 and 4 days (mean ± SEM). **d)** Maximal F/F_0_ of PtR1 cells grown in P_limited_ conditions with 1.8 µM phosphate for four days following resupply by different concentrations of phosphate (n.b. cells were grown on NSW, but artificial seawater (ASW) was used for the phosphate resupply experiments to abolish additive effects from ambient phosphate in NSW). Prior to resupply cells were pre-perfused for 15 s with ASW medium without phosphate or other nutrients). Cells (*n*) examined over 3 independent experiments (mean ± SEM). **e**) Maximal F/F_0_ of PtR1 cells grown in P_limited_ conditions with 1.8 µM phosphate for four days following resupply with 36 µM of different phosphorus forms (e), including: phosphate (Pi), adenine triphosphate (ATP), glucose 6-phosphate (G6P), polyphosphate (PolyP), or bis(*p-* nitrophenyl)phosphate (b-NPP). Cells (*n*) examined over 3 independent experiments (mean ± SEM).

Ambient phosphate concentrations can vary significantly in coastal waters. Levels at L4 station in the Western English Channel, where diatom blooms are seen frequently, can reach ∼0.8 µM in February/March to lower than 0.05 µM in July (Archer et al., 2009). Transitory spikes up to 0.97 µM during summer phosphate concentration minima have also been reported (Jordan and Joint, 1998), providing phosphate resupply opportunities in phosphorus-limited phytoplankton populations. To determine the sensitivity of the phosphate-Ca^2+^ signalling response, we carried out a dose-response experiment. Exposure of four-day P_limited_ cells to resupply revealed that cells responded to environmentally-relevant phosphate concentrations as low as 0.9 µM (**Figure 1d**). Our control condition (artificial seawater (ASW) without phosphate), did not induce a response. The described phosphate-Ca^2+^ signalling pathway thus exhibits high sensitivity to inorganic phosphate concentrations within the range of those seen in natural ecosystems, and is therefore of broad environmental relevance.

Phosphorus in the oceans can exist in numerous forms. This includes both inorganic (e.g. phosphate and polyphosphate) and organic forms. Dissolved organic phosphorus (DOP) can exceed orthophosphate concentrations (Björkman and Karl, 2003; Karl and Bjorkman, 2002), with phosphoesters often the dominant class (Kolowith et al., 2007). We tested the efficacy of different phosphorus forms for activating the Ca^2+^-signalling response in four-day P_limited_ PtR1 cells. Treatment with equimolar concentrations (36 µM) of phosphomonoesters (adenosine triphosphate, ATP and glucose-6-phosphate, G6P), or inorganic polyP all led to transient elevations in cytosolic Ca^2+^, similar to those evoked by phosphate (**Figure 1e**). In contrast, the phosphodiester bis(*p-*nitrophenyl)-phosphate (b-NPP) did not. We found that *P. tricornutum* can grow unimpaired on all of the different phosphorus forms examined, albeit at a significantly reduced specific growth rate with b-NPP (**Figure S1d**). These results indicate that exposure of P_limited_ cells to phosphorus forms besides phosphate (with the exception of b-NPP) can evoke [Ca^2+^]_cyt_ elevations. However, it is unclear whether this is because the phosphorus-Ca^2+^ signalling pathway can perceive these forms directly, or whether phosphoesterases convert them to inorganic phosphate prior to detection. Extracellular ATP is also a well-known signalling molecule in plants and animals, which can trigger Ca^2+^-dependent signalling pathways regardless of phosphorus status (Tanaka et al., 2014). We therefore tested the efficacy of these compounds on P_replete_ cells, since we have established that phosphate-Ca^2+^ signalling is activated only in phosphate-limited cells (**Figure 1b**). We did not detect Ca^2+^ elevations in response to any of these phosphorus forms in P_replete_ cells (**Figure S1e**). Moreover, treatment of 4-day P_limited_ cells with a poorly hydrolysable form of ATP, ATPγS (Tang et al., 2003), did not yield [Ca^2+^]_cyt_ elevations (**Figure S1f**). Although these results do not exclude the possibility that different phosphorus forms can directly trigger the phosphorus-Ca^2+^ signalling pathway, they strongly suggest that phosphate-stressed *P. tricornutum* cells can rapidly liberate phosphate from organic phosphorus forms (likely via extracellular phosphatases upregulated during phosphorus stress (Cruz de Carvalho et al., 2016; Diaz et al., 2018; Lin et al., 2013)), which subsequently evoke a Ca^2+^ response. This is further supported by our evidence that b-NPP did not evoke a [Ca^2+^]_cyt_ elevation. Hydrolysis rates are reportedly considerably slower for b-NPP than for phosphomonoester substrates in phosphorus-limited *P. tricornutum* cells (Flynn et al., 1986). Thus longer-term processes appear to be necessary to liberate b-NPP, as is also supported by the reduced growth rate of *P. tricornutum* on this substrate (**Figure S1d**). Taken together, these data indicate that phosphorus-Ca^2+^ signalling can be evoked, albeit indirectly, via a range of environmentally abundant phosphorus forms.

### Rapid crosstalk between P and N metabolism following phosphate resupply revealed by comparative proteomics and stable-isotope tracer experiments

To determine how the phosphorus-Ca^2+^ signalling pathway may regulate cellular adaptations to phosphate amendment, we employed a comparative proteomics approach to determine early recovery responses from phosphorus limitation. We grew PtR1 cells in three regimes, including P_replete_, P_limited_, and P_resupply_ treatments for four days. We then resupplied 36 µM to P_limited_ cells (for the P_resupply_ treatment) and harvested all cultures four hours later. Total proteins were then extracted for comparative proteomics analysis. From the 1505 identified proteins (**Dataset S1**), 443 were differentially expressed (exhibiting a log2 fold change ≥ 1; ≤-1, Q < 0.05) in P_limited_ versus P_replete_ cells (215 were more abundant, and 228 were less abundant). By comparison, 232 proteins had significantly altered relative abundance in P_resupply_ versus P_limited_ cells (63 increased and 169 decreased abundance) (**Figure S2 Dataset S2-S3**). We classified differentially expressed proteins into specific metabolic pathways, using Mercator-based analyses (Lohse et al., 2014). This identified broad-scale impacts of phosphate regime on proteins associated with protein, nitrogen, DNA and RNA metabolism, cell division, alongside photosynthesis and signalling (**Figure S2; Dataset S2-S3**), consistent with previous studies (Cruz de Carvalho et al., 2016; Wurch et al., 2011).

We also observed significant enhancement of phosphate acquisition and recycling proteins in P_limited_ versus P_replete_ cells (Alipanah et al., 2018; Cruz de Carvalho et al., 2016). Indeed, under P limitation, the four most highly-expressed proteins included a glycerophosphoryl diester phosphodiesterase and three predicted alkaline phosphatases (**Dataset S3**). These proteins remained highly expressed 4 h following phosphate resupply. By comparison, notably, some of the most significantly altered proteins in the P_resupply_ treatment (compared to P_limited_ cells) related to nitrogen uptake/assimilation. This included upregulation of a nitrate transporter (NRT1) that showed a striking 5.7 log2 fold increase, alongside five other nitrogen metabolism proteins (**Figure 2a and b**). Glutamate dehydrogenase (GD) and chloroplast-targeted fd-glutamate synthase (fd-GOGAT) also exhibited decreased abundance. These data suggest that a major immediate response to phosphate resupply in diatoms is the upregulation of nitrogen uptake. However, further experimental characterisation is necessary to determine the timescale and impact of such changes on actual nitrogen uptake rates, how they are regulated, and their importance to cell physiology.

**Figure 2.**
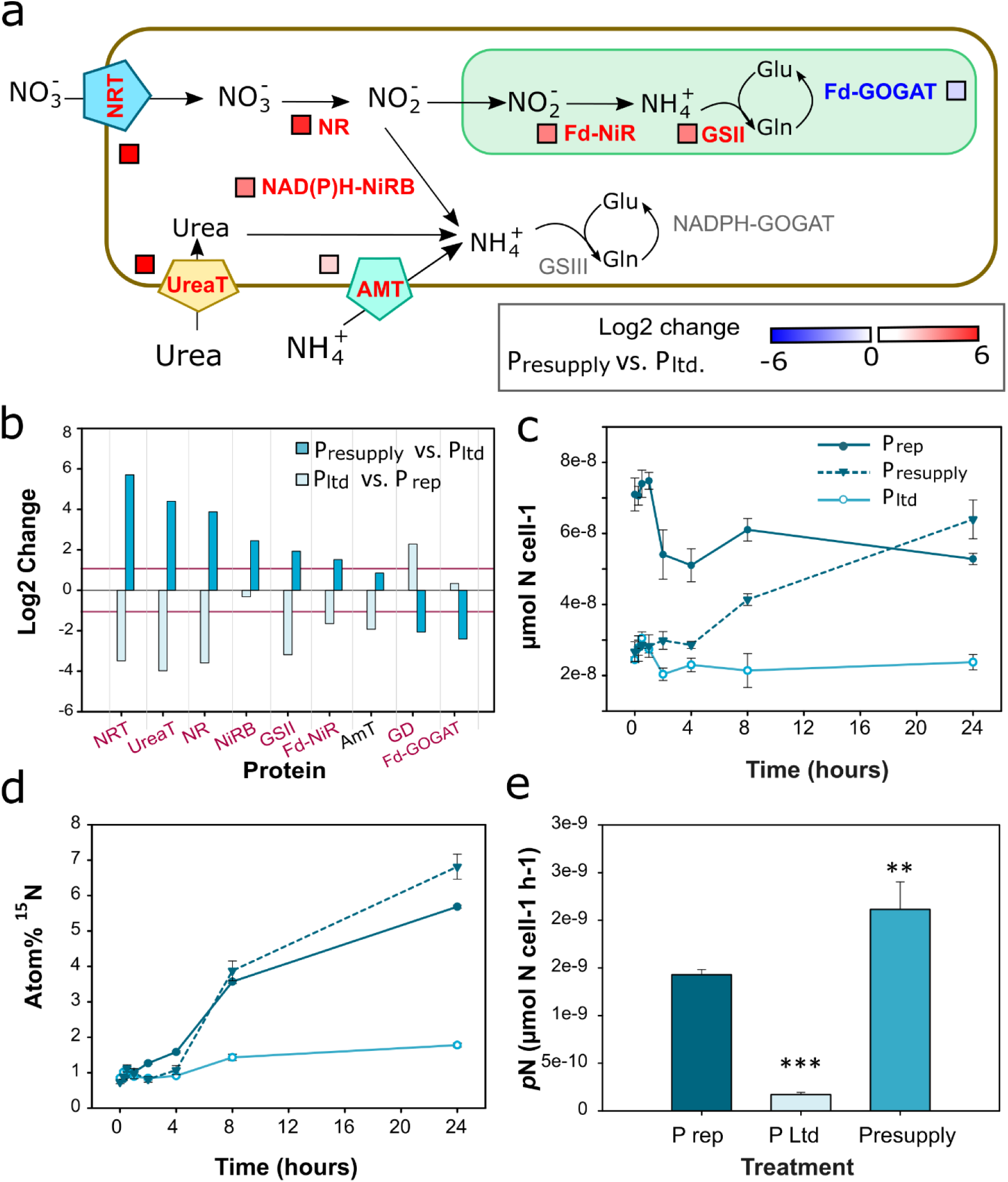
Rapid crosstalk between P and N metabolism following phosphate resupply. **a)** A cohort of proteins associated with nitrogen uptake and assimilation exhibited altered abundance after four hours in P_resupply_ compared to P_limited_ cells. This included increased abundance of a nitrate transporter (NRT/NRT2; JGI protein identifier: 26029/54101), a urea transporter (20424/768), nitrate reductase (NR; 54983), NAD(P)H-dependent nitrite reductase (NirB; 13154), chloroplast-targeted glutamine synthetase (GSII; 51092), and ferridoxin-dependent nitrite reductase (NiR; 12902). We also saw decreased abundance of a putative glutamate dehydrogenase (45239), and the chloroplast-targeted glutamate synthase (GOGAT/GltS, 56605). **b)** Bar graph of protein fold changes of nitrogen metabolism proteins described in (a) in response to phosphate resupply (relative to P_limited_ cells; log2 fold change ≥ 1, Q < 0.05 labelled purple). The log2 fold changes of proteins exhibiting significantly altered abundance in P_limited_ cells relative P_replete_ cells are also shown. **c-d**. total nitrogen uptake (µmol N cell^-1^) (c) and atom% ^15^N (d) in P_replete_, P_limited_ and P_resupply_ treatments following phosphate resupply to phosphate-limited cells over 24 hours (mean (*n*=3) ± SEM). **e**) Absolute nitrate uptake rates (pN) µmolN cell^-1^ h^-1^ of P_replete_, P_limited_ and P_resupply_ cultures following phosphate resupply to phosphate-limited cells over 24 h (mean (*n*=3) ± SEM). Asterisks (*) indicate statistically significant differences (one-way ANOVA, ***p<0.001, ** p<0.01) compared to the phosphate-replete control.

To directly examine the impact of phosphorus resupply on nitrogen uptake over time, we characterised changes in total cellular nitrogen content, and ^15^N-nitrate uptake, in PtR1 cells experiencing different phosphate regimes. We grew up P_replete_, P_limited_ and P_resupply_ treatments for four days. Prior to phosphate resupply we added ^15^N-nitrate (to a concentration 10% of ambient nitrate) to all the cultures and acclimated cells for one hour. We then quantified the total nitrogen content and ^15^N enrichment (expressed as atom% ^15^N) over 24 hours following phosphate resupply. At T_0_ (i.e. just prior to phosphate resupply), P_replete_ cells had 2.9 times more total nitrogen than P_limited_ cells (**Figure 2c**). However, upon phosphate resupply significant increases in total nitrogen content were detected within just eight hours, and levels exceeding those in P_replete_ cells were measured in 24 hours. By comparison, the cellular nitrogen content of P_limited_ cells remained low. Moreover, approximately nine-fold increases in atom% ^15^N levels occurred within 24 hours following phosphate resupply (**Figure 2d**). By comparison, the levels in P_limited_ cells did not increase beyond initial values. Moreover, absolute nitrate uptake rates were twelve times greater in P_resupply_ compared to P_limited_ cultures, and 1.5 times more than the P_replete_ cells over 24 hours (**Figure 2e**). These data demonstrate that the proteomic changes observed in the abundance of nitrogen transport proteins, as a consequence of P resupply, result in rapid and substantial increases in nitrogen uptake.

### Enhanced nitrogen transport is a primary acclimation response driving recovery from phosphate stress

We have observed enhanced nitrogen transport in P_limited_ cells within just eight hours of phosphate amendment. Nitrogen is a major constituent of proteins, nucleic acids, chlorophyll, and other macromolecules. Alongside proteomic changes in nitrate transport machinery, we observed concomitant increases in numerous proteins of protein metabolism in P_resupply_ versus P_limited_ cells. This included the increased abundance of 19 synthesis proteins and concomitant decreased abundance of 13 degradation proteins (**Figure S2; Dataset S2**). Examination of the total protein content of cells grown under different phosphate regimes, confirmed levels were significantly reduced in P_limited_ compared to P_replete_ cells, but subsequently recovered following phosphate resupply after 24 hours, prior to increases in cell density (**Figure 3a-b**). In addition, we saw altered abundance of 13 photosynthesis-related proteins in P_resupply_ versus P_limited_ cells. Nine of these were down-regulated, and they all related to the light reactions of photosynthesis, including five light harvesting complex (LHC) proteins, Cytochrome B6 (PetB), alongside two proteins of photosystem II (PsbC and PsbA) (**Dataset S2)**. These changes correlated with photo-physiological adaptations in P_resupply_ cells, including increases in Fv/Fm values, and total chlorophyll within 24 hours (**Figure 3c-d**). Additionally, a rapid reduction in non-photochemical quenching (NPQ) was seen six hours following phosphate resupply (**Figure 3e**) that was not detected in P_limited_ or P_replete_ cells (**Figure 3f**). These NPQ changes were on a time-frame more similar to those observed for nitrogen transport (**Figure 2c-d**). NPQ is a vital photo-protection mechanism for dissipating harmful excess light energy. Notably, the observed reductions in NPQ caused increases in overall photosynthetic capacity of P_resupply_ cells, as demonstrated by increased electron transport rates (ETR µmol e m^-2^ s^-1^) (**Figure 3e, red**). This will likely enhance photosynthetic reducing power, which could potentially drive other processes such as nitrate assimilation. Phosphate-driven adaptations in NPQ have also been reported in *Micromonas*, and are hypothesised to occur via an ancient light-harvesting-related protein, LHCSR (LHCX proteins in diatoms) (Guo et al., 2018). We detected significant upregulation of one LHCX protein (Lhcx3) in P_limited_ cells (compared to P_replete_ conditions) (**Dataset S3**). This protein is involved in light-driven adjustments in NPQ in *P. tricornutum* (Buck et al., 2019). Our data thus suggests Lhcx3 may also contribute to NPQ adaptations to phosphate stress in diatoms.

**Figure 3.**
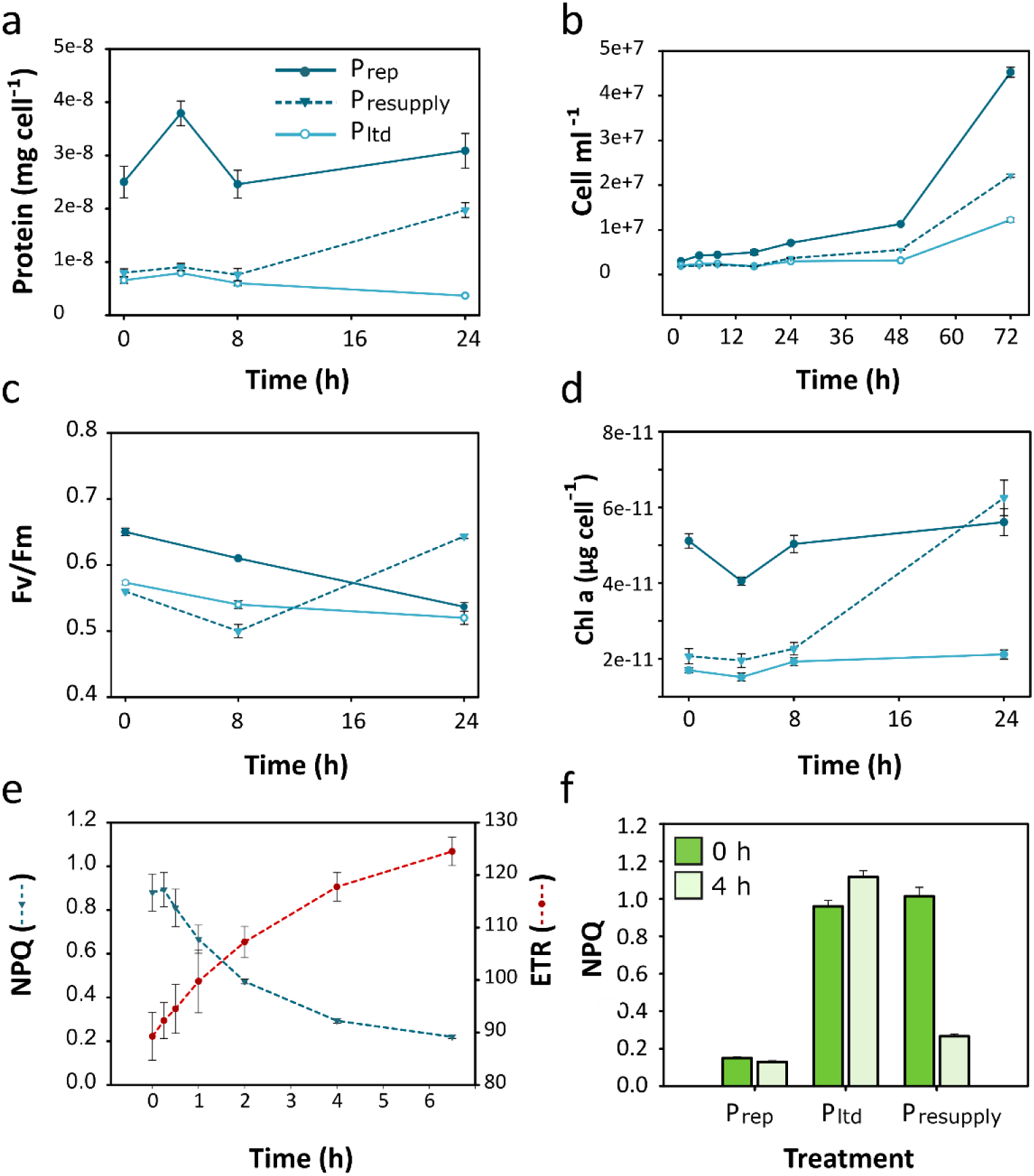
A succession of metabolic acclimation responses drive recovery from phosphate stress. Changes over time of **a)** Total protein content (mg cell^-1^), **b**) cell density (cells ml^-1^), **c**) Fv/Fm, and **d**) total chlorophyll a (chl a) content (µg cell^-1^) of phosphate-limited (P_limited_; 1.8 µM) cultures following resupply (P_resupply_) of phosphate (36 µM), compared to P_replete_ (36 µM) and P_limited_ grown cells. **e**) Changes in non-photochemical quenching (NPQ) and electron transport rate (ETR, µmol electrons m^-2^ s^-^1) in P_limited_ cells in response to phosphate resupply over six hours, **f**) NPQ values of P_replete_, P_limited_, and P_resupply_ prior to (0 h) and four hours (4 h) after phosphate resupply.

A key response of diatoms for coping with P stress is the accumulation of neutral lipids including triacylglycerides (TAG) and replacement of membrane phospholipids with non-P forms (Van Mooy et al., 2009). However, we detected just five proteins of lipid synthesis/metabolism exhibiting differential abundance (log2 fold change ≥ 1, Q < 0.05) in P_resupply_ versus P_limited_ cells (**Dataset S2**). This included the decreased abundance of one enzyme of phospholipid metabolism (cyclopropane-fatty-acyl-phospholipid synthase), alongside two fatty-acid biosynthesis enzymes: enoyl-CoA hydratase (ECH1) and acetyl-CoA carboxylase (ACC1), and concomitant increased abundance of a fatty acid desaturase. ECH1 and ACC1 catalyse the synthesis of precursors for TAG biosynthesis (Balamurugan et al., 2017). The downregulation of these proteins corresponds with the recovery of TAG levels to those similar to P_replete_ cells, 24 hours following phosphate resupply (**Figure S3**).

Together, our evidence points to a metabolic succession of recovery responses from phosphorus stress following phosphate resupply. This begins with substantial increases in nitrogen uptake alongside NPQ adjustments (and increases in ETR) within just 8 hours, and is followed by recovery of protein, TAG and chlorophyll levels, Fv/Fm and subsequently growth. The primary adaptations of nitrogen metabolism observed thus likely underpin subsequent cellular adaptations necessary to kick-start cellular growth following phosphate amendment.

### Phosphate-Ca^2+^ signalling is necessary for primary adaptations in nitrate metabolism following phosphate resupply

We have characterised early physiological adaptations underpinning phosphate stress recovery, and identified an important role for nitrogen uptake and NPQ adaptation, within hours of phosphate resupply. Alongside acclimation of primary metabolism, many proteins associated with cell signalling were regulated by phosphate. Notably, this included the abundance of numerous Ca^2+^-signalling-related proteins in P_limited_ versus P_replete_ cells, including several Ca^2+^/calmodulin-dependent protein kinases (**Dataset S3**; **Figure S4**) that could serve as sensors for phosphorus-induced Ca^2+^ elevations, as has been documented in the N-Ca^2+^ signalling response of *Arabidopsis* (Liu et al., 2017). Several of these were also downregulated following phosphate resupply (**Dataset S2)**. These findings uncover putative mechanistic components of the pathway, and add further evidence to the importance of Ca^2+^-signalling in phosphorus sensing, and likely role in regulating metabolic adaptations to phosphorus resupply.

To examine whether the phosphate-Ca^2+^ signalling pathway mediates downstream recovery responses from P limitation following phosphate resupply, we investigated the source and pharmacology of the phosphate-induced Ca^2+^ signal with the aim to identify avenues to inhibit phosphate-Ca^2+^ signals. Treatment of P_limited_ cells to phosphate resupply in artificial seawater lacking Ca^2+^ (+ 200 µM EGTA) completely abolished the phosphate-induced Ca^2+^ elevation (**Figure 4a**), indicating dependency of the response on external Ca^2+^. This suggests that plasma membrane localised channels are involved. *P. tricornutum* encodes a number of Ca^2+^ channel homologues (Verret et al., 2010), for which there has been little/no functional characterisation. This is with the exception of a novel class of channels that we recently characterised in diatoms (EukCatAs) (Helliwell et al., 2019). We therefore examined whether *Pteukcata1* knockout lines (Helliwell et al., 2019) are impaired in phosphate-Ca^2+^ signalling. All lines tested evoked phosphate-induced [Ca^2+^]_cyt_ elevations comparable to PtR1 (**Figure 4b**), indicating that EukCatA1 is not involved in the primary Ca^2+^ response to phosphate. We also adopted a pharmacological approach, testing the effect of Ca^2+^ channel blockers on phosphate-Ca^2+^ signals. These experiments revealed that whereas pre-treatment of cells with verapamil (L-type Ca^2+^ channel inhibitor) did not disrupt the phosphate-Ca^2+^ signal (**Figure S5a**), 5 µM Ruthenium Red (RuR; inhibits a range of Ca^2+^ channels (Vincent and Duncton, 2011) inhibited the response (**Figure 4c**). By comparison, 5 µM RuR did not disrupt Ca^2+^ signalling responses to hypo-osmotic stress, which is known to trigger cytosolic Ca^2+^ elevations in *P. tricornutum* (Falciatore et al., 2000; Vardi et al., 2006) (**Figure S5b & c**). Thus RuR does not interfere with the capacity of R-GECO1 to report Ca^2+^, or cause broad disruption of Ca^2+^ signalling processes within the cell.

**Figure 4.**
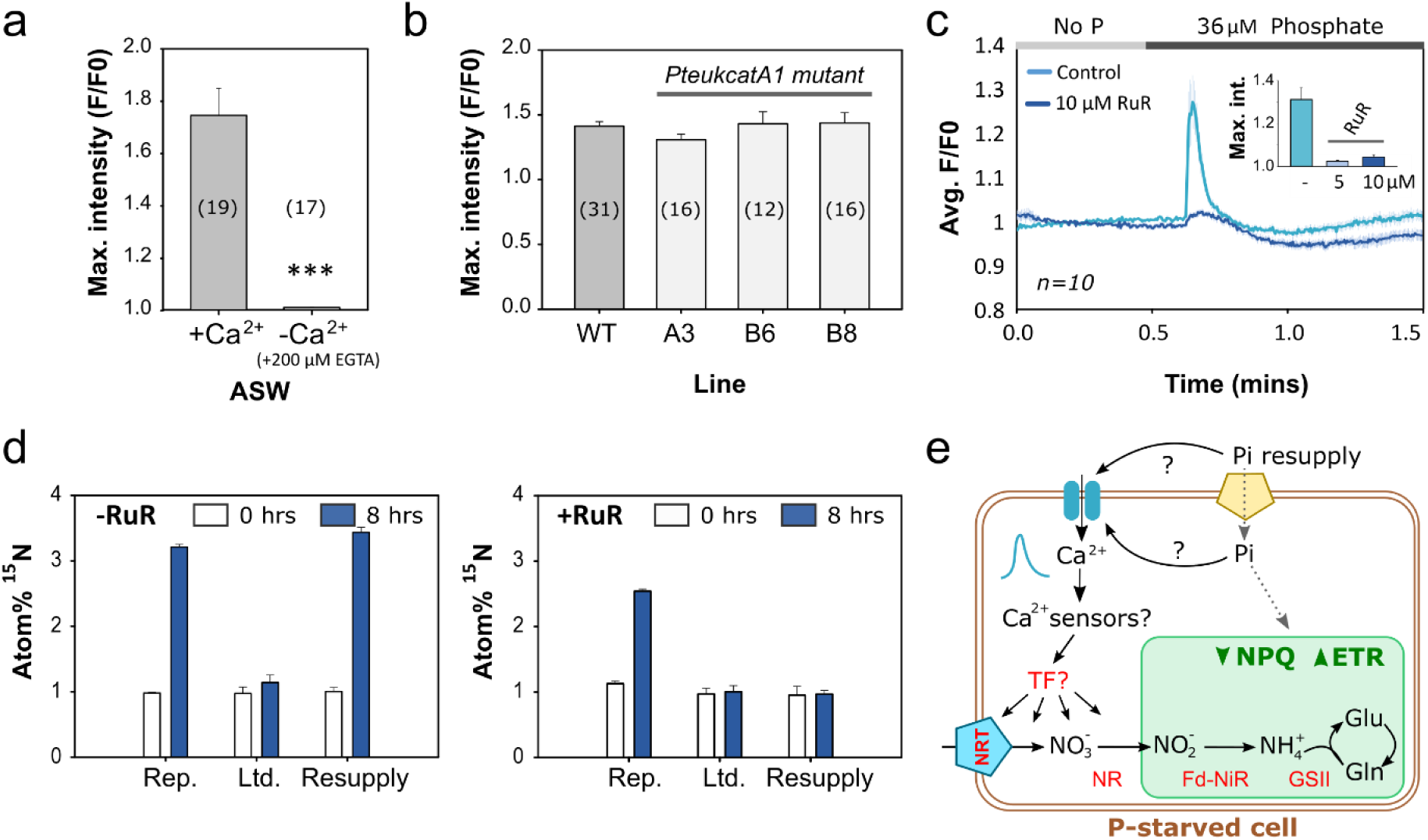
Phosphate-Ca^2+^ signalling is necessary for nitrate uptake following phosphate resupply. **a)** Average maximal fluorescence (F/F0) of four-day PtR1 cells grown in artificial seawater (ASW) with limited phosphate (1.8 µM) exposed to phosphate resupply (36 µM) either with or without Ca^2+^ (+ 200 µM EGTA). No. of cells (*n*) examined over 3 independent experiments each with a different sample of cells (mean ± SEM), *p*-value (students t-test) ***, <0.001). **b**) Comparison of average maximal fluorescence (F/F0) values of phosphate-Ca^2+^ signalling response in PtR1 cells versus three independent *PteukcatA1* mutant lines in a PtR1 background (Helliwell et al., 2019). Cells were grown in standard f/2 medium (with natural seawater, NSW), with low phosphate (1.8 µM) prior to the experiment. Cells (*n*) were examined over 3 independent experiments per line each with a different sample of cells (mean ± SEM). **c**) Average fluorescence trace of the phosphate-Ca^2+^ signalling response in PtR1 cells pre-treated for 5 minutes with 10 µM RuR, compared to control (no inhibitor). Prior to phosphate resupply cells were pre-perfused for 30 s with NSW medium without RuR, phosphate or other nutrients. Inset displays average maximal F/F0 values of the phosphate-Ca^2+^ signalling response without inhibition (-), and following treatment with 5 µM RuR, 10 µM RuR (*n*= 17, 19 and 13 cells over three independent experiments; mean ± SEM). **d**) Atom% ^15^N measured in cells under different phosphate regimes in the absence or presence of 5 µM RuR. The atom% ^15^N measured at T=0 (prior to phosphate resupply), and 8 hours post phosphate resupply is shown (*n*=3; mean ± SEM; experiments were repeated twice with similar results). **e**. Schematic model for the novel phosphorus-Ca^2+^ signalling pathway. Ca^2+^ independent components are indicated with dashed arrows (key: TF: transcription factor, ETR: electron-transport rate, NPQ: non-photochemical quenching, Pi: phosphate, NRT: nitrate transporter, NR: nitrate reductase, fd-NiR: ferridoxin-dependent nitrite reductase, and GSII: glutamine synthetase.

We therefore investigated the effect of inhibiting the phosphate-Ca^2+^ signalling pathway on early primary adaptations (i.e. nitrogen uptake and NPQ) to phosphate amendment in P_limited_ cells. We first examined the impact of RuR inhibition on the nitrate uptake response, limiting exposure of cells to RuR to 8 hours (i.e. when enhanced nitrate uptake was first detectable in P_resupply_ cells, Figure 2d). As observed previously, we detected a significant increase in atom% ^15^N following P resupply within 8 hours, which was absent in P-limited cells (**Figure 4d, left**). Similar increases in atom% ^15^N were also observed in the P_replete_ treatments in the presence or absence of RuR (5 µM) (**Figure 4d**), indicating that nitrate uptake is not inhibited by RuR. By comparison, in the P_resupply_ treatment +5 µM RuR, no increases in atom% ^15^N levels were observed at all. To examine whether the altered nitrate uptake in the P_resupply_ +RuR treatment was due to reduced cell health following incubation with RuR, we measured Fv/Fm values. However, exposure of P_resupply_ cells to 5 µM RuR for 8 hours did not reduce Fv/Fm values, compared to the –RuR control (**Figure S5d**). Finally, we tested the impact of RuR on NPQ following phosphate resupply. Whereas NPQ values were not perturbed by four hour treatments with RuR in P_replete_ and P_limited_ cells (**Figure S5e**), the fast reductions in NPQ following phosphate resupply over six hours were still observed following phosphate resupply (**Figure S5f**). Therefore, RuR treated P_limited_ cells still exhibit phosphate-induced NPQ recovery responses.

Together, our findings demonstrate that the rapid changes in NPQ capacity can occur in a Ca^2+^-independent manner, potentially responding directly to increased cellular P quotas detectable within hours following phosphate resupply in *P. tricornutum* (Cáceres et al., 2019). By comparison, fundamental increases in nitrate uptake in P_limited_ cells following phosphate resupply are dependent on phosphate-induced [Ca^2+^]_cyt_ elevations (**Figure 4e**). Thus the P-Ca^2+^-signalling pathway is vital for regulating primary metabolic recovery from P limitation, and also serves to maximise acquisition and resource competition for the vital limiting nutrient N.

## Discussion

We report the discovery of a novel Ca^2+^ signalling pathway in diatoms to sense and rapidly respond to increases in phosphorus availability (**Figure 4e**). We show that the addition of phosphorus to phosphate-limited *Phaeodactylum* cells results in cytosolic Ca^2+^ elevations within seconds of resupply. This response is evoked by environmentally-relevant phosphate concentrations, and (indirectly) by different phosphorus forms. Moreover, inhibition of phosphate-Ca^2+^ signalling completely blocks a critical component of the recovery response from phosphate limitation (nitrate uptake) that underpins subsequent physiological responses. Phosphorus-Ca^2+^ signalling is therefore likely a critical driver of competitive phytoplankton dynamics, and important to survival in dynamic nutrient environments, such as upwelling, coastal and estuarine systems where phosphate concentrations vary significantly (Jordan and Joint, 1998). More broadly, these findings provide much needed insight into the molecular mechanisms employed by eukaryotic algae for sensing phosphorus, which until now have remained enigmatic (Dyhrman, 2016).

Our work also highlights that fundamental cross-talk between the essential nutrients phosphorus and nitrogen drive ecological acclimation to phosphorus availability in diatoms. Phosphorus-limited cells invest primarily in phosphate acquisition (Alipanah et al., 2018; Cruz de Carvalho et al., 2016), scavenging (Dyhrman et al., 2012; Ly et al., 2014) and reallocation (Martin et al., 2011), diverting resources away from vital processes such as nitrogen assimilation. Meanwhile, activation of the phosphate-Ca^2+^ signalling machinery readies cells for detection of phosphorus resupply, which governs rapid phosphate-driven induction of nitrogen-assimilation proteins. This allows phosphorus-limited cells to control the allocation of resources to priority cellular functions, which must then be rapidly rewired when conditions change. This rapid coordination between P and N metabolism, mediated via the phosphorus-Ca^2+^ signalling pathway, enables diatoms to immediately exploit another vital limiting nutrient once released from phosphorus limitation. As phosphate resupply events (e.g. due to riverine inputs/heavy rainfall, upwelling, or microscale cell lysis processes) (Litchman, 2007) often occur simultaneously with enhanced nitrate abundance (Smyth et al., 2010), by upregulating nitrogen assimilation, the phosphorus-Ca^2+^ signalling pathway primes the cell to anticipate improved nutrient conditions more generally. Notably, this cohort of nitrogen-related genes are key indicators of phytoplankton nitrogen status (McCarthy et al., 2017), and can also exhibit rapid responsiveness to nitrate resupply (Smith et al., 2019). However, we found no evidence for a role of Ca^2+^ signalling in nitrate sensing in N-limited *P. tricornutum* cells. Together these data highlight that multiple environmental drivers coordinate resource-responsive gene expression in diatoms, via complex regulatory networks.

Finally, this study expands the portfolio of biological functions of Ca^2+^ signalling known. Diatoms are evolutionarily divergent from plants and animals, in which Ca^2+^ signalling research is well established. By broadening our study to important taxa outside of ‘crown’ eukaryote groups, we can gain a much more comprehensive understanding of the role and evolution of Ca^2+^ signalling across the eukaryote tree of life. Indeed, by taking this approach, we have identified that distinct mechanisms for nutrient perception have arisen in photosynthetic eukaryotes. Diatom-like phosphorus-Ca^2+^ signalling is apparently absent in plants: phosphate-induced [Ca^2+^]_cyt_ elevations were not detected in phosphorus-stressed *Arabidopsis* (Matthus et al., 2019). In a similar vein, unlike *Arabidopsis*, nitrate resupply did not evoke a Ca^2+^-signalling response in N-limited *Phaeodactylum* cells. Nevertheless, the diatom phosphate-Ca^2+^ signalling pathway does share features with Ca^2+^-dependent nitrate-sensing in *Arabidopsis* (Liu et al., 2017). Both pathways coordinate expression of nitrogen-related genes via Ca^2+^. In *Arabidopsis*, this is orchestrated by Ca^2+^ sensor-kinases that phosphorylate NIN-LIKE PROTEIN (NLP) transcription factors. Intriguingly, NLP transcription factors are absent from diatom genomes (Rayko et al., 2010). However, we did find four Ca^2+^ sensor-kinase genes upregulated during phosphorus limitation. Notably, they also contain recognition motifs for the phosphorus-stress Myb-like transcriptional regulator PtPSR (Sharma et al., 2019). Our work thus paves the way to future advances in our understanding of the genetic components, evolutionary distribution, and broader roles of phosphate-Ca^2+^ signalling in controlling phosphorus-stress recovery in diatoms, and potentially eukaryotes more broadly.

## Materials and Methods

### Cultivation of *P. tricornutum*

*Phaeodactylum tricornutum* strain CCAP1055/1 was obtained from the Culture Collection of Algae and Protozoa (SAMS limited, Scottish Marine Institute, Oban, UK). The transgenic line R-GECO1(Zhao et al., 2011) (PtR1) was generated as described by Helliwell et al., (2019) (Helliwell et al., 2019). Cultures were maintained in natural seawater (NSW) supplemented with f/2 nutrients (Guillard and Ryther, 1962), with 100 μM Na_2_SiO_3_.5H_2_O (but not vitamins) unless stated otherwise, and illuminated with 50-80 µmol m^-2^ s^−1^ light, on a 16∶8 h light:dark cycle at 18°C. For experiments with artificial seawater (ASW), the following recipe was used: 450 mM NaCl, 30 mM MgCl_2_, 16 mM MgSO_4_, 8 mM KCl, 10 mM CaCl_2_, 2 mM NaHCO_3_, and 97µM H_3_BO_3_), with f/2 nutrients + Si. For all physiology and signalling experiments *P. tricornutum* cultures were inoculated to a cell density of 3 × 10^4^ cells ml^-1^ in liquid culture.

For the nutrient (N, P, f/2 metals) limitation treatments for nutrient resupply experiments described in **Figure S1a**, cells were grown in f/2 medium made up with NSW, but with concentrations of nitrate or phosphate reduced to one twentieth of those typically found in standard f/2 medium (Guillard and Ryther, 1962), and no f/2 metals for the trace metal limitation treatment (Met). Cell densities in these different nutrient conditions were: 2.8 × 10^6^ cells ml^-1^ (nutrient replete), 1.6 × 10^6^ cells ml^-1^ (P limited), 1.6 × 10^6^ cells ml^-1^ (N limited) and 4 × 10^5^ cells ml^-1^ (met limited), on the day of the nutrient resupply experiments.

### Epifluorescence imaging in *P. tricornutum*

*P. tricornutum* cells grown in liquid culture for 4 days were placed in a 35 mm glass-bottomed dish (In Vitro Scientific, Sunnyvale, CA, USA) coated with 0.01% poly-L-lysine (Sigma-Aldrich, St Louis, MO, USA). Cells adhered to the bottom of the dish were imaged at 20°C using epifluorescence microscopy with a Nikon Eclipse Ti microscope with a 100×, 1.30 NA oil immersion objective and detection with a Photometrics Evolve EM-CCD camera (Photometrics, Tucson, AZ, USA). Excitation of R-GECO1 (PtR1) cells was performed using a pE2 excitation system (CoolLED, Andover, UK) with 530-555 nm excitation and 575-630 nm emission filters. Images were captured using NIS-ELEMENTS v.3.1 software (Nikon, Japan) with a 300 ms camera exposure (frame rate of 3.33 frames s^−1^). Images were processed using NIS-ELEMENTS v.3.1 software. The mean fluorescence intensity within a region of interest over time was measured around the perimeter of each cell. Change in fluorescence intensity of R-GECO1 was calculated by normalising each trace by the initial value (F/F0).

During imaging, cells were continuously perfused with natural seawater (3 ml min^−1^). The phosphorus resupply treatments were delivered by switching the perfusion from f/2 medium without phosphate to f/2 medium with phosphate (typically 36 µM, except in **Figure 1d** for the phosphate dose experiment), unless otherwise stated. Cells exposed to phosphate resupply treatments in the absence of Ca^2+^ were perfused with at least 20 ml Ca^2+^ free medium (+200 µM EGTA) in order to minimise residual Ca^2+^ from the ASW medium. The same set-up was used for hypo-osmotic shock experiments, except the perfusion was switched from undiluted to diluted NSW.

### Treatment with pharmacological inhibitors for Ca^2+^-signalling experiments

Prior to phosphate resupply and/or hypo-osmotic shock treatments, cells were bathed in seawater containing verapamil (5 µM), Ruthenium Red (RuR, 5 µM or 10 µM) for 5 minutes in glass-bottomed dishes (stock solutions for these chemicals were made up in ddH_2_O). Experimental treatments switching from medium containing no phosphate to 36 µM phosphate (or to diluted seawater in the case of the hypo-osmotic shock experiments) without the pharmacological agents were then delivered, as outlined above.

### Protein preparation for shotgun proteomics

P_replete_, P_limited_, and P_resupply_ treatment cultures were inoculated with PtR1 cells to a cell density of 3 × 10^4^ cells ml^-1^ and incubated in standard growth conditions for 4 days. We added 36 µM of phosphate to the P_resupply_ cultures and incubated all cultures for an additional 4 h at 18°C. Cells from 15 ml of culture were harvested by centrifugation (4000 g at 4°C for 5 min). Supernatants were removed and cell pellets flash-frozen until further analysis. Cell pellets were then dissolved in 100 µl of 1 x LDS loading buffer (Invitrogen, USA) and given three cycles of 5 min sonications (Branson 2510 Ultrasonic water bath), 10 s of vortex and 5 min incubations at 95°C. Thirty µl of the lysate was loaded immediately onto a precast 10% Tris-Bis NuPAGE gel (Invitrogen, USA) using 1 x MOPS solution (Invitrogen, USA) as the running buffer. SDS-PAGE was performed for a short gel migration (5 mm of migration into the gel). Polyacrylamide gel bands containing the cellular proteomes were excised and standard in-gel reduction with dithiothreitol and alkylation with iodoacetamide was performed prior to trypsin (Roche, Switzerland) proteolysis (Christie-Oleza and Armengaud, 2010). The resulting tryptic peptide mixture was extracted from the polyacrylamide gel bands and prepared for mass spectrometry as previously described (Kaur et al., 2018).

### NanoLC-MS/MS and data analysis of the proteomes

Samples were analysed by nanoLC-ESI-MS/MS using an Ultimate 2000 LC system (120 minute LC separation on a 25 cm column; Dionex-LC Packings) coupled to an Orbitrap Fusion mass spectrometer (Thermo-Scientific, USA), using LC conditions and MS settings as described previously (Christie-Oleza et al., 2015). Raw MS/MS files were processed with MaxQuant version 1.5.3.30 (Cox et al 2008) for protein identification and quantification, using default parameters, match between runs and *P. tricornutum* strain CCAP 1055/1 protein database (Ref. UP000000759) obtained from UniProt. The comparative proteomic analysis between samples (i.e. data filtering and processing, as well as two-sample Student’s T-tests and fold changes) was carried out using Perseus version 1.5.5.3 (Tyanova et al., 2016) following the pipeline described previously (Kaur et al., 2018), but including a stringent rule where only proteins confidently detected in all three biological replicates of at least one condition were considered. The full list of detected proteins is available in Dataset S1.

### Biochemical analyses

#### Total protein extraction and quantification

For total protein analyses, 2 mL of cells were spun down for 2 mins at 13,000 g at 20°C, the supernatant removed, and pellets flash-frozen in liquid nitrogen. Cell pellets were then re-suspended in 50-200 µL (according to original cell density) of protein extraction buffer (comprising of 2% SDS, 5 mM tris-HCL pH 6.8, cOmplete™ protease inhibitor cocktail (1 tablet per 50 ml of extraction buffer), and sonicated for 3 mins in a sonication bath with ice. Total protein was then quantified using a Pierce BCA Protein Assay kit (Thermo Fisher Scientific), according to manufacturer’s instructions.

#### Chlorophyll quantification

To measure total chlorophyll concentrations a 2 mL aliquot of *P. tricornutum* cells was centrifuged for 2 mins at 13,000 g at 21°C, the supernatant discarded and cell pelleted re-suspended in 1 ml ethanol. Chlorophyll pigments were extracted by vortexing for 2 mins, followed by centrifugation at 13,000 g for 2 mins at 21°C. Optical density of the supernatant was then measured at 652, 665, and 750 nm, and the equations from Ritchie et al., (2008)(Ritchie, 2008) applied to calculate chlorophyll *a* concentration per cell.

#### Neutral lipid staining and quantification

One mL of cells were stained with 1 µl of 25 µg ml^-1^ Nile Red dissolved in DMSO. Fluorescence was then measured in a CLARIOstar plate reader (BMG LABTECH) using the AlexaFluor532 pre-setting (excitation/emission settings: 482±16/570-530).

### Photo-physiological measurements

Measurements of Fv/Fm and non-photochemical quenching (NPQ) were made on dark-adapted (15 minutes) cells using an AquaPen-C device (Photon Systems Instruments). For NPQ measurements, different treatments were diluted to equivalent cell densities prior to quantification (OD_730_ between 0.02-0.03). The predefined NPQ3 setting was used (light duration 200s, 10 pulses; dark recovery duration 60s, 2 pulses), with light intensity settings as follows: 450 µmol.m^-2^.s^-1^ (actinic light), 3000 µmol.m^-2^.s^-1^ (super-pulse i.e 100%) and 20% for the flash pulse. Values for Φ_PSII_ (QY_LSS_) were then extracted to calculate electron transport rate (ETR) using the following equation: Φ_PSII_ × photosynthetically active radiation (PAR) × 0.5 (where PAR was 450 μmol photon m^−2^ s^−1^ (actinic light)), according to Maxwell et al., (2000) (Maxwell and Johnson, 2000).

### Nitrogen uptake experiments

P_replete_, P_limited_, and P_resupply_ treatment cultures were inoculated with PtR1 cells to a cell density of 3 × 10^4^ cells ml^-1^ and incubated in standard growth conditions for 4 days. Sodium nitrate-15N (≥98 atom% ^15^N, ≥99%, Sigma Aldrich, 364606) was added to the cultures (to a concentration 10% of the ambient nitrate) (Dugdale and Goering, 1967), and cells incubated for 1 h at 18°C. Phosphate (36 µM) was then added to the P_resupply_ cultures. Cells were harvested at each time-point by centrifugation at 4000 g for 10 mins at 4°C, supernatant removed and pellets snap-frozen in liquid nitrogen. Analysis of particulate nitrogen and atom %15N were made by continuous flow mass spectroscopy (Rees et al., 1999).

## Acknowledgments

We acknowledge support from the NERC-IRF grants NE/R015449/2 (K. H), NE/K009044/1 (J. C-O), ERC grant ERC-ADG-670390 (C. B.), and the NERC-CLASS programme grant (NE/R015953/1; A. R). We are also grateful to the Proteomics Research Technology Platform (University of Warwick, UK), and Malcolm Woodward (PML, UK) for experimental support.

## Author Contributions

K.H, G.W and A.R designed the experiments; K.H., E.H, J.D, J.C-O, M.A-F and L.A-M conducted the experiments. K.H., A.R., and J.C-O. analysed the data. K.H. wrote the paper with input from G.W, J.C-O, C.B., and A.R.

## Supplementary Videos

**Video S1**. Ca^2+^ imaging of 4-day old phosphorus-limited *P. tricornutum* (PtR1) cells, following resupply with 36 µM phosphate. Cells were pre-perfused with standard f/2 medium (made up natural seawater) without phosphate for 30 secs prior to perfusion with f/2 medium with 36 µM phosphate.

## Supplementary Figures

**Figure S1.**
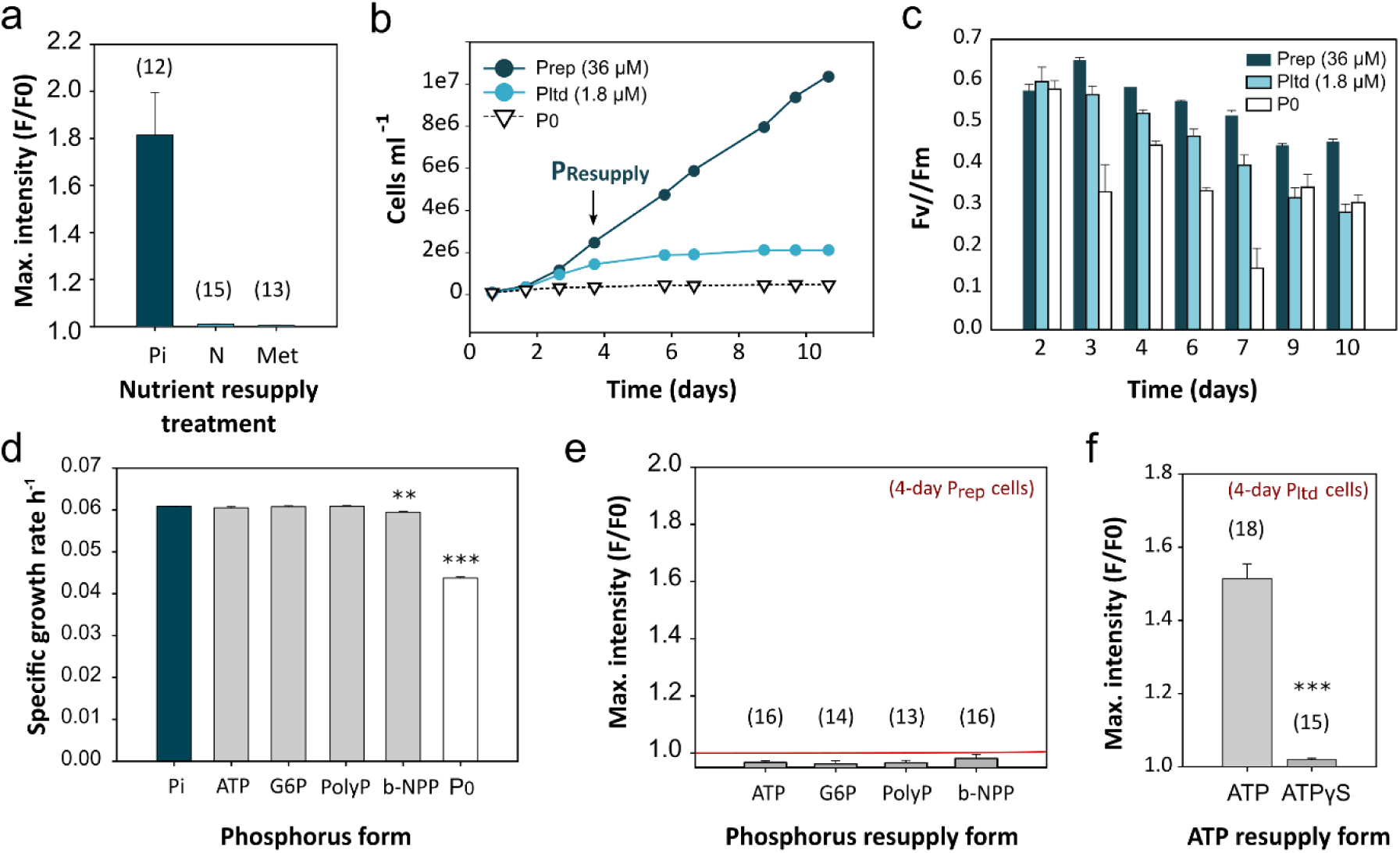
Novel phosphorus-Ca^2+^ signalling mechanism is activated in phosphate-limited cells by a range of phosphorus forms. **a)** Mean maximal fluorescence (F/F_0_) of PtR1 cells grown for four days with limiting concentrations of either phosphate (P), nitrate (N) or metals (Met), exposed to natural seawater (NSW) with phosphate, nitrate or Met restored to full f/2 concentrations (Guillard and Ryther, 1962). For the nutrient limitation treatments, cells were grown in f/2 medium made up with NSW, but with concentrations of nitrate or phosphate reduced to one twentieth of those typically found in standard f/2 medium (Guillard and Ryther, 1962), or no f/2 metals for the trace metal limitation treatment (Met). Cell densities in these different nutrient conditions were: 2.8 × 10^6^ cells ml^-1^ (nutrient replete), 1.6 × 10^6^ cells ml^-1^ (P limited), 1.6 × 10^6^ cells ml^-1^ (N limited) and 4 × 10^5^ cells ml^-1^ (met limited), on the day of the nutrient resupply experiments. Cells were pre-perfused with seawater for 30 secs prior to nutrient amendments. Number (*n*) of cells examined over 3 independent replicate experiments carried out with a different sample of cells for each replicate is shown; error bars represent SEM. **b**) Growth over time of PtR1 cells in standard f/2 medium with phosphate-replete (P_replete_, 36 µM), phosphate-limited (P_limited_, 1.8 µM) or no phosphate amendment (P_0_) conditions (*n*=3; SEM). Arrow indicates the time at which phosphate was resupplied to P_limited_ cells for the P_resupply_ treatments described in this study. **c**) Quantum yield (Fv/Fm) of cells at different time-points over the course of the experiment for the growth curve displayed in (b) (*n*=3; SEM). **d**) Specific growth rate (h^-1^) of PtR1 cells grown in f/2 medium with 36 µM phosphate (Pi), ATP, G6P, PolyP or b-NPP as a phosphorus source (*n*=3; SEM). Asterisks (*) indicate statistically significant differences (ANOVA, \*\*\**p*<0.001, ** *p*<0.01) compared to the phosphate control. **e)** Mean maximal fluorescence (F/F_0_) of PtR1 cells grown for four days in standard f/2 medium (i.e. phosphate-replete conditions) in response to f/2 medium without inorganic phosphate, but amended with 36 µM adenosine triphosphate (ATP), glucose-6-phosphate (G6P), polyphosphate (PolyP), or bis-(*p*-nitrophenyl)phosphate (b-NPP). Cells were pre-perfused with standard f/2 (natural seawater) medium without phosphate for 30 secs prior to perfusion with f/2 medium (including 36 µM of the phosphorus form being tested). Number (*n*) of cells examined over 3 independent replicate experiments is shown; error bars represent SEM. **f)** Comparison of maximal fluorescence (F/F0) response of four-day old, P_limited_ cells in response to 36 µM ATP versus the poorly-hydrolysable ATPγS form (Tang et al., 2003). Number (*n*) of cells examined over 3 independent replicate experiments is shown using a different sample of cells for each replicate; error bars represent SEM (*p* value (Student’s ttest): ***, <0.001).

**Figure S2.**
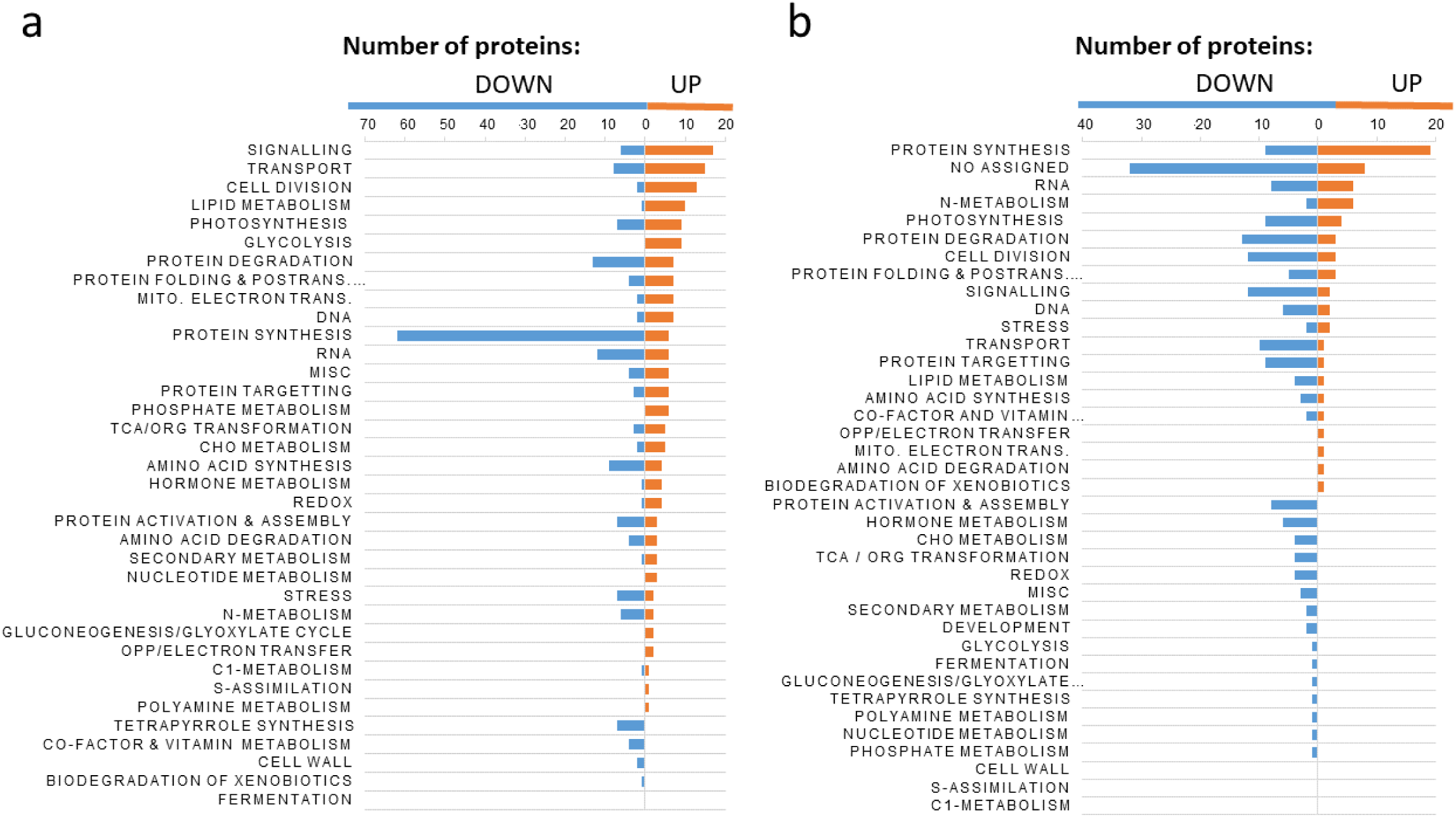
Overview of annotated functions of *Phaeodactylum tricornutum* proteins with significantly altered abundance in cells grown under different phosphate regimes. **a**. Proteins with increased (up) and decreased (down) abundance in PtR1 cells grown for four days in low phosphate (P_limited_; 1.8 µM) compared to replete (P_replete_; 36 µM) conditions (of proteins showing a log2 fold change ≥ 1, Q < 0.05; *n*=3). Proteins are categorized into major functional groups based on Mercator analysis (Lohse et al., 2014). **b**. As in a), but for P_resupply_ versus P_limited_ cells. Cells from all treatments (P_replete_, P_limited_ and P_resupply_) were harvested four h following resupply of 36 µM of phosphate to the P_resupply_ treatment.

**Figure S3.**
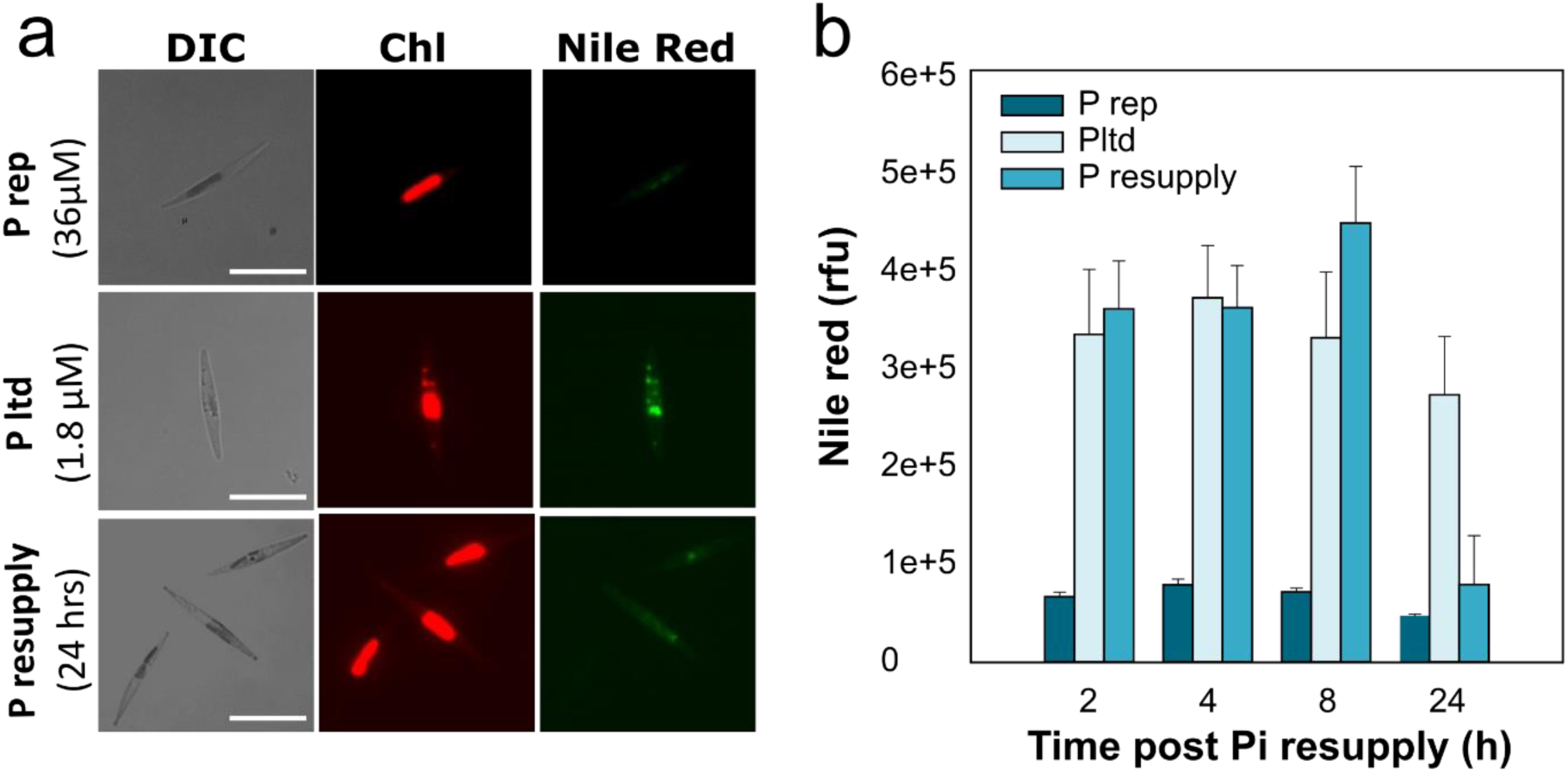
Quantification of lipid bodies of *P. tricornutum* cells in phosphate-replete, limited and resupply treatments. **a**. Epifluorescence image of WT *P. tricornutum* cells grown for four days in i) P_replete_, ii) P_limited_ and iii) P_resupply_ cells stained with Nile Red to visualise lipid bodies (green). Chlorophyll auto-fluorescence (red) and differential interference contrast (DIC) images are also shown. Scale bar: 10 µm. **b**) Average fluorescence intensities (methods) of P_resupply_ cells over time following amendment with 36 µM phosphate, compared to P_replete_ and P_limited_ cells (that had no phosphate resupply).

**Figure S4.**
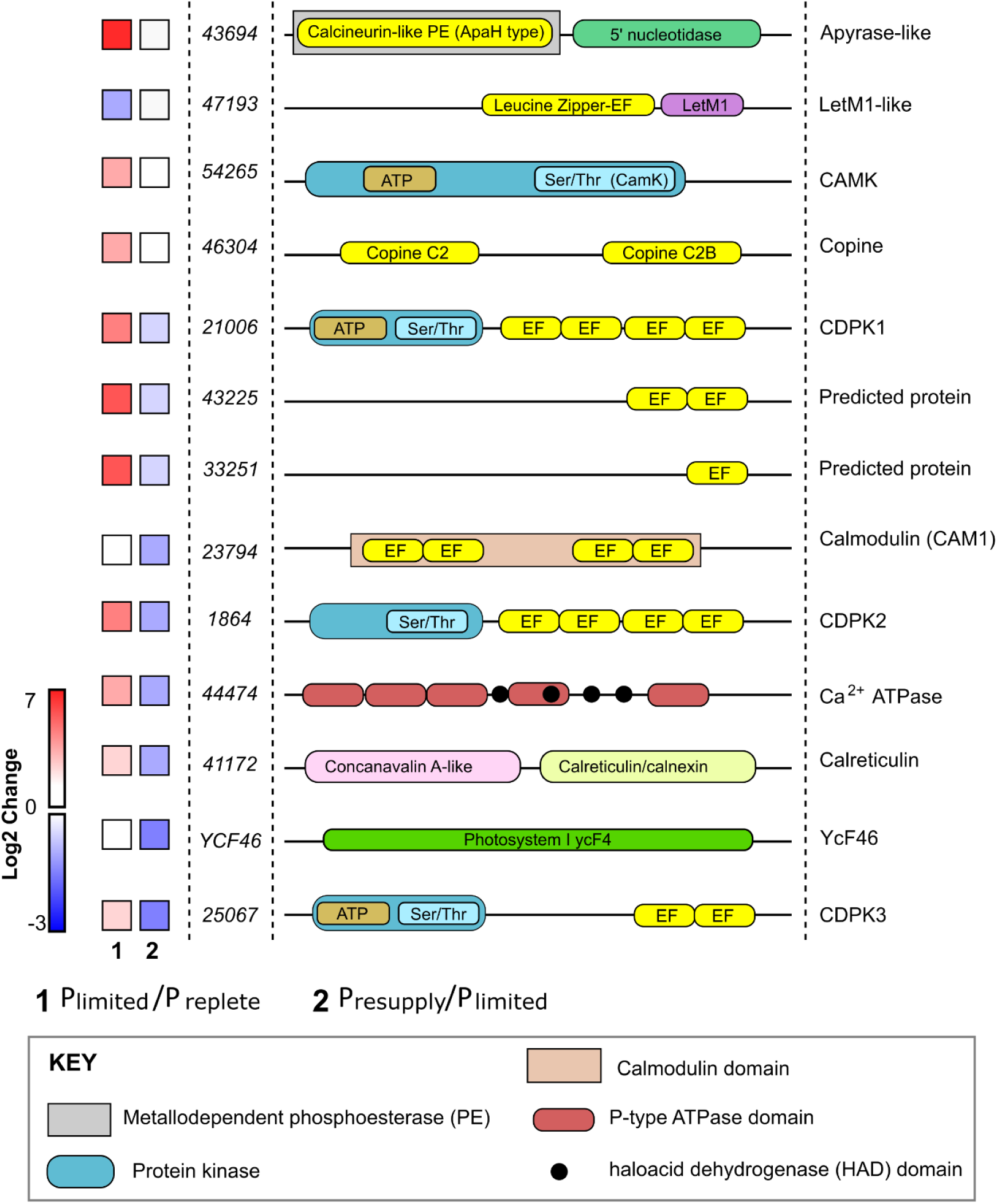
Overview of Ca^2+^-signalling-associated proteins altered in abundance by phosphate. A schematic diagram of the domain structure of thirteen *P. tricornutum* proteins assigned to Ca^2+^-signalling functions that had significantly altered abundance (showing a log2 fold change ≥ 1, Q < 0.05; *n*=3) in P_limited_ compared to P_replete_ (1) samples and P_resupply_ compared to P_limited_ (2) treatments. The proteins described and their associated JGI protein identifiers are as follows: apyrase-like protein (apyrase-L; 43694), mitochondrial proton/calcium exchanger protein (LETM-L; 47193), Ca^2+^/calmodulin-dependent protein kinase (CAMK2; 54265), Copine (46304), Ca^2+^-dependent protein kinase (CDPK1; 21006), EF-hand containing predicted proteins (43225 and 33251), Calmodulin (CAM1; 23794), CDPK2 (1864), ATPase1-2B (44474), calreticulin (CALR; 41172), YCF46 (Uniprot identifier: A0T0G7), and CDPK3 (25067). Domains analyses were conducted using Interpro (Apweiler et al., 2000).

**Figure S5.**
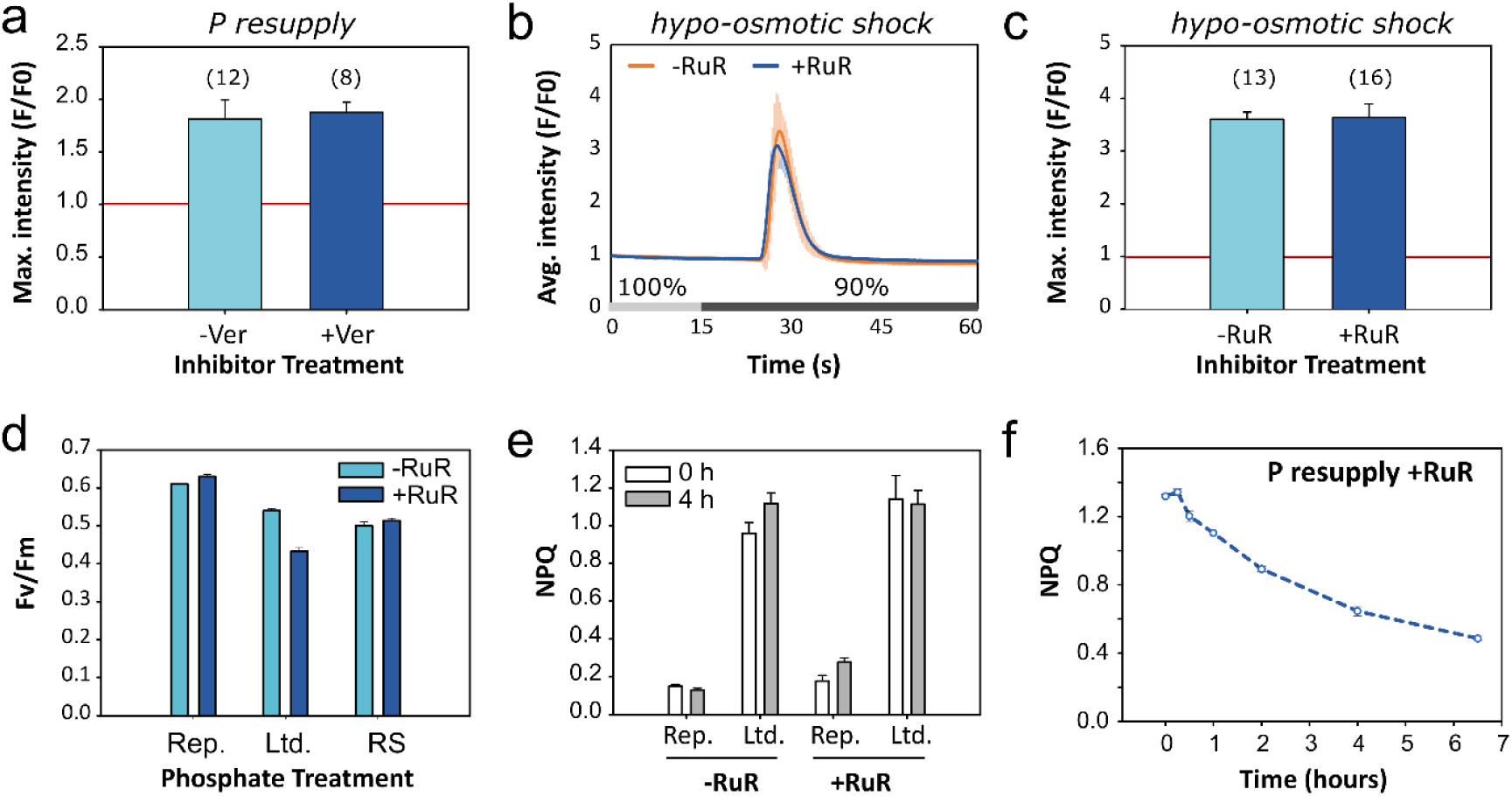
Characterising the effect of pharmacological inhibiters on phosphate-Ca^2+^ signalling. **a)** Ca^2+^ channel blocker verapamil does not affect phosphate-induced Ca^2+^ signals. Maximal amplitude (F/F0) of phosphate-limited cells (grown for four days on f/2 medium with 1.8 µM phosphate), and pre-exposed to 5 µM verapamil (+Ver) following resupply with 36 µM phosphate, compared to the –Ver control. Prior to resupply cells were pre-perfused for 15 s with natural sweater (NSW) without Ver, phosphate or other nutrients. Number of cells examined over 3 independent replicate experiments each with a different sample of cells is shown in parentheses; error bars represent SEM. **b & c**) Treatment of PtR1 cells ruthenium red (RuR; 5 µM) does not impair hypo-osmotic shock induced Ca^2+^ signalling. Four-day old PtR1 cells were pre-treated with 5 µM RuR for five minutes, and then perfused for 15 s with 100% NSW (without RuR, or nutrients), before being exposed to 90% NSW (90% seawater; 10% deionised water). Average fluorescence (F/F0) traces (b) and maximal fluorescence (c) following exposure to the hypo-osmotic shock are plotted. Number of cells examined over 3 independent replicate experiments each with a different sample of cells is shown in parentheses; error bars represent SEM. **d**) Fv/Fm values of four-day old P_replete_, P_limited_ and P_resupply_ treatments following 8 h incubation with 5 µM RuR, compared to the –RuR control (*n*=3; SEM). **e**) NPQ values of four-day old P_replete_, and P_limited_ treatments following four h incubation with 5 µM RuR, compared to the –RuR control (*n*=3; SEM). NPQ values prior to RuR treatments (i.e. 0 hrs) are also shown (*n*=3; SDM). **f**) NPQ of low phosphate grown WT *P. tricornutum* cells treated with 10 µM RuR, in response to phosphate resupply (*n*=3; error bars represent SEM).

